# Open source computational simulation for a moth-inspired navigation algorithm

**DOI:** 10.1101/744912

**Authors:** Noam Benelli, Roi Gurka, Yiftach Golov, Ally Harari, Gregory Zilman, Alex Liberzon

## Abstract

Olfactory navigation in insects, for instance when males search for mates, is a navigational problem of a self-propelled agent with limited sensor capabilities in a scalar field (odor) convected and diffused by turbulent wind. There are numerous navigation strategies proposed to explain the navigation paths of insects to food (flowers) or mating partners (females). In a search for a mate, the males use airborne pheromone puffs in turbulent environments around trees and vegetation. It is difficult to compare the various strategies because of a lack of a single simulation framework that can change a single parameter in time and test all the strategies against a controlled environment. This work aims at closing this gap, suggesting an open source, freely accessible simulation framework, abbreviated MothPy. We implement the simulation framework using another open source package (“pompy”) that recreates a state-of-the-art puff-based odor plume model of Farrell et al. [1]. We add four different navigation strategies to the simulation framework based on and extending the previously published models [2, 3], and compare their performance with different wind and odor spread parameters. We test a sensitivity analysis of the navigation strategies to the plume meandering and to increased turbulence levels that are effectively expressed as the elevated puff spread rates. The simulations are compared statistically and provide an interesting view on the robustness and effectiveness of various strategies. This benchmarking-ready simulation framework could be useful for the biology-oriented, as well as engineering-oriented studies, assisting to deduce the evolutionary efficient strategies and improving self-propelled autonomous systems in complex environments.

## Introduction

The known ability of male moths to reach conspecific females from long distances (hundreds to thousands of meters) in turbulent environments such as forests and canopies, [4, 5] has attracted considerable interest in both biological and engineering studies devoted to navigation strategies, e.g. [2, 3, 6, 7]. Moths are able to navigate efficiently using only local cues mediated by turbulent air. This is accomplished by using chemo-receptors on their antennae [8–10] for chemical sensing [11, 12] whilst using visual optometry for the spatial orientation, using so-called *optomotor anemotaxis*. The navigational paths of moths were mostly observed to perform relative narrow zigzagging [13–15] motion and wider side-slips, sometimes called “casting” or “sweeping” motion, respectively. A large variety of models were proposed to explain their navigation strategy, using an internal counter [16–19] or a different set of assumptions [13, 20–24].

In addition, there are probabilistic types of navigation models that use the olfactory signal with or without prior information or memory assumptions, e.g. [2, 6, 23, 25–30]. In order to evaluate the feasibility, accuracy and readiness of the current odor-based navigational models, there is a need for a framework that will enable an assessment of the variability of the proposed models. A unified framework and a computer simulation for quantitative comparison of the above mentioned models and others, similar to those proposed in autonomous navigation studies [1, 29–31] is required.

Recently, Macedo et al. [32] reported about a simulator and comparison of several bio-inspired and engineered strategies for chemical plume tracking. However, this framework is based on the diffusion process without accounting for wind or turbulence that are at the core of the moth-inspired navigation strategies [2, 3, 6, 29]. In this work, we provide a framework using an open source computational platform with wind and plume characteristics set as parameters that can be adjusted in order to more realistically simulate the environmental conditions.

Herein we examine few available moth-inspired navigation strategies based on a prescribed wind and plume model. The main goal is to provide an accessible and reproducible scientific simulation platform, promoting the development of new navigation strategies for biology and autonomous vehicles.

## Methods and materials

We build a benchmarking and comparison framework based on numerical simulations for odor-based navigation. We utilize a well-known wind and plume model of [1] provided by an open source software package [33] to which we add navigation models, as described below. We provide the framework as an open source package written in Python and entitled MothPy, see [34].

In the following section we briefly review the wind and plume models, described in details in [1] and with greater details, a set of four *navigation models* chosen for this work. The models are different by the concept and their construction. Therefore, we have chosen a common set of sub-modules. For instance, if some models include both casting or foraging parts, and other models do not have such parts, we compare only the same parts (casting/zigzagging/surging) . The choice is rather arbitrary, yet has a common base that enables a fair comparison.

### Computational framework

The computational framework is an open source package written in Python [34]. It is based on the open source scientific software packages of Numpy, Scipy, Matplotlib and Jupyter, among others. The core of the wind and plume model was developed by another open source project, [33], and adopted here as a good representation of the model of Farrell et al. [1]. The software package is developed as the object oriented programming concept, containing several classes for the wind, plume and basic navigator properties. The four examples of navigators were implemented as objects from those classes for the present case study. The perspective user can easily install the required packages using standard Python procedures and develop its own classes based on these examples. For easier adoption and reproducibility, we developed an online cloud-based Jupyter notebook (use the link from the software repository) in the spirit of a new term in the scientific world *preproducibility* [35]

### Wind and plume model

The simulation is performed in a numerical domain representing a two dimensional flow field in the streamwise - spanwise plane. It resembles a horizontal plane parallel to the ground at some height above the vegetation without obstacles. The two velocity components are the streamwise or wind direction component, *u*, and the transverse or cross-wind direction, *v* in the direction of *y* axis. Generally, the crosswind component is about an order of magnitude weaker than the streamwise component, but not negligible.

In some cases, a user might want to create a more realistic representation of turbulent plumes in complex environments. The wind model contains an option to include random noise which can represent turbulent velocity fluctuations [1]. We added also a periodic, large-amplitude and relatively slow (relative in respect to the flight time of a navigator) component that mimics the meandering of the plume. For simplicity, both the random and the periodic components are added to all the grid nodes of the flow field, without any spatial variation along and across the simulation field. This setup does not replicate features of real turbulent flows, yet it creates an effectively similar, turbulent-like, chemical plume, because as time propagates and puffs move, every part of the plume experiences only local changes in velocity. Although this wind-plume model is not a physical model, its strength is in providing a reasonably fast simulation framework for testing multiple navigators and obtaining a convergent statistics.

Mathematical formulation of the wind model is a wind vector field prescribed at every location *x*, *y* as:

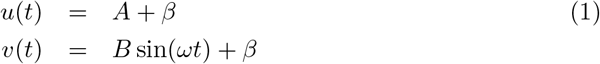

where *A*, *B*, *ω* are constants, chosen as the simulation parameters of the wind speed, meandering amplitude and period, respectively. Parameter *β* represents a random white noise. For more detailed technical information on the implementation of the wind model, see [1].

### Plume model

The plume model simulates the release of odor from a point source located upstream in respect to the navigator. We call it odor, although the released scalar represents equally temperature or concentration, among others [1]. Conceptually, the source emits the odor through so-called “puffs”. Physically speaking, a puff is a finite amount of scalar that is preserved in time during its dispersion (a mass conservation). Mathematically, it can be defined as a two-dimensional Gaussian shape carried downstream by the wind, defined by the coordinates of the center of the puff, *x*_*p*_(*t*), *y*_*p*_(*t*) and the concentration distribution around the puff *C*, which is related to the standard deviation of the Gaussian function. The puff center, *x*_*p*_(*t*), *y*_*p*_(*t*), will move in the flow field with a speed determined by the two-dimensional wind vector (*u*, *v*). For simplicity, we use bold mathematical symbols for the 2D vectors, i.e. **u**= {*u*, *v*}, **x**_*p*_ = {*x*_*p*_, *y*_*p*_}:

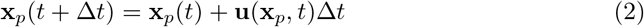

The concentration *C*(*r*, *t*) around a single puff is determined by the distance from the puff’s center, **x**_*p*_(*t*), as well as the time passed since the formation of the puff *t*.

Broadly speaking, as the puff “matures” it becomes more dispersed, in a two-dimensional approximation mathematically described in Eq. (3):

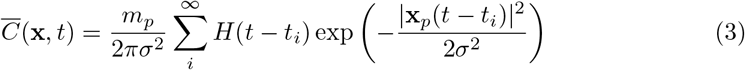

where *m*_*p*_ is the mass of the puff, *H* is a Heaviside function and *σ* is the spreading rate proportional to turbulent diffusivity, see e.g. Ref. [3] for the references therein. In the following, we simplify the problem using the agents with a binary sensor, therefore the concentration of odor is translated into the size of the region in which the concentration is above a threshold of detection, i.e. *C* ≥ *C*_0_. The size in this approximate model is a circular patch of radius *r*_*p*_(*t*) and its area is proportional to *r*_*p*_(*t*)^2^, growing at the rate proportional to *σ*^2^:

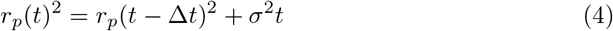

Parameters of the source are the puff release rate, *f*_*r*_ (in units of frequency, puffs per second) and puff spread rate, *dr*_*p*_(*t*)*/dt*. The spread rate defines the linear rate of increase of *r*_*p*_(*t*), as shown in Eq. (4). These two source parameters, together with the wind parameters and the concentration threshold of the agent’s odor sensor, determine the characteristic of the plume. For instance, setting the threshold to a negligible value will convert the plume type from an array of discrete, concentrated puffs into a single, featureless stream of odor. We are interested in the present case study in a downwind spreading plume of odor, resembling in some sense a trail of puffs, similar to that made by the conditions in a wind tunnel with a single female moth secreting pheromone [36].

### Navigation strategies

We chose four navigation strategies for the present case study, tested in the proposed computational framework. These are based on Ref. [3] (named here as: “A”,“B”), and on Ref. [2] (named here as: “C”,“D”). An overview of each navigational strategy is provided in the following.

#### Definitions

The strategy of the navigator model is comprised from a set of rules and constrains that underline the decision making process. For the cases studied here, there are several assumptions that are similar for all the different strategies:

- The agent is a free-flying object travelling at a constant ground speed and a binary sensor (yes/no) for the odor cues.
- The agent can only measure the local wind direction and it can use an internal counter [16] for the time scale estimates.
- The agent does not have a long-term memory or spatial information in respect to the fixed coordinate system (no GPS signal)

Mathematically, we describe the agent as an object marked by a point in a two dimensional space, 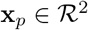, a point-sensor of the local wind velocity, **u**(**x**), and presence/absence of odor *c*(**x**_**p**_) = 1*/*0. Although a flying navigator will only sense wind velocity relative to itself, we assume that using optometry data, the navigator can find the direction of the wind relative to the ground. Here we adopt the widely acceptable notion of *optomotor anemotaxis* [2, 29]. That assumption is in accordance with the directly observed behaviour of moths in a wind tunnel and previous models [1, 3].

Navigation in our study is performed in a simulated wind and concentration field, as explained above. The binary sensor threshold of a navigator is the last parameter that defines the field for a given navigator. Two such examples of the identical simulation fields for the two navigators with two different thresholds are shown in Fig. 1. In this figure, regions that would be tagged “detected” puffs are marked by white pixels and the background (the concentration below the threshold) is dark. It is noteworthy to mention that physically identical plumes (same wind, turbulence, release rate, concentration) may appear very different to different navigators depending on their detection threshold, as in Fig. 1. We address this issue in the discussion.

**Fig 1.**
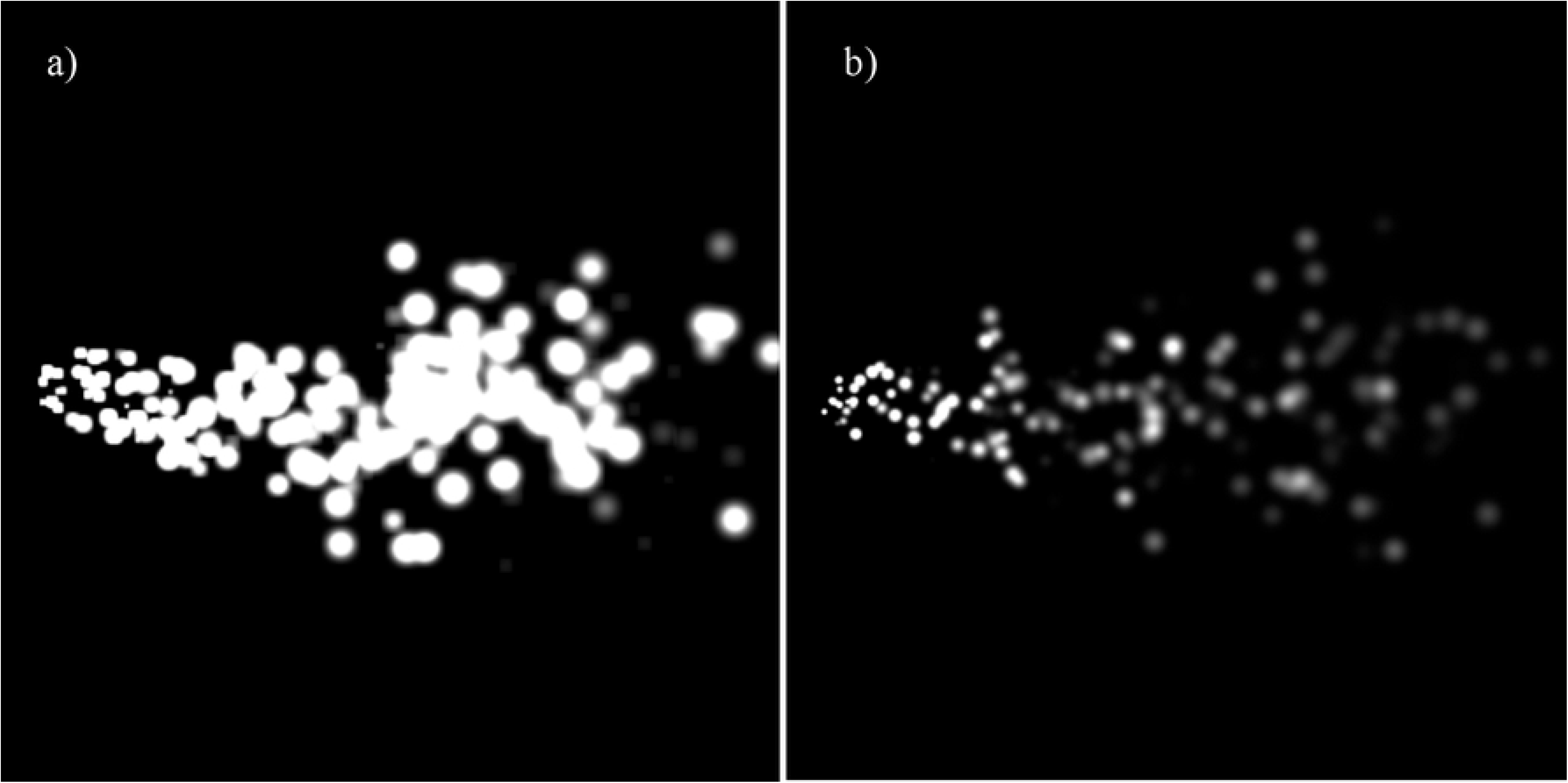
Two examples of the plumes with discrete puffs, based on the model of [1], using [33]. The wind direction is from left to right and the source (“female moth”) is at the origin located at the center of the left side of the figure. The plume release parameters and wind parameters are identical, and the difference in the visualized puffs is due to the threshold limit of the “male moth” binary sensor: a) 1500 and b) 30000 (arbitrary units). Concentration above a threshold is marked by white pixels.

We focus on the navigation part and not on the complete process of foraging or mating search. Therefore, in our simulations, the flyer does not actively search for the plume. The initial condition is that a navigator is placed at position *x*_0_, *y*_0_ downstream the source (*x*_0_ *>* 0) and within an area with a certain probability to encounter a puff. The navigation starts when a puff with concentration above a given threshold “touches” the initial location of the navigator. This moment is marked as the initial time of the navigation path *t*_0_. The navigation path consists of several possible time intervals:

- “detection” - the time of flight during which the navigator is inside a puff, i.e. the measured concentration is above the threshold;
- “surging” - straight upwind flight after detection interval;
- “casting” or “zigzagging” - crosswind flight with alternating changes of direction, typically when the signal has been recently lost or uncertain;
- “sweeping” - large random motions that are designed to increase the probability to encounter a next signal.

These elements of the navigation strategy are similar to those observed in moth flights, see e.g. [2].

#### Strategy “A”

Strategy “A” is based on the navigational strategy developed in Ref. [3]. This strategy is based on an essential parameter: the puff crossing time, or in the terms of our definitions, “detection time”. This time will be denoted as *t*_*c*_ and it is reset every time a navigator crosses a puff. We will use the same notation for all the navigation strategies. After detection, surging (fast upwind motion according to the local wind direction) will be set for some time, called λ, see Fig. 2. In some cases, the time λ will be a constant (predefined time interval), however, in navigator “A” it is determined by the previous detection time, λ = *t*_*c*_. For the strategy “A”, a navigator that does not meet a new puff within the time of surging λ will start casting with the transverse zigzags of time intervals, denoted here as *δ*_1_. Similarly to λ, *δ*_1_ can be a predefined constant, or a variable. In this particular navigator “A”, the time interval *δ*_1_ is like the surging time interval, proportional to the last detection or crossing time *t*_*c*_ as it follows λ = *αt*_*c*_. In the particular case shown in the results *α* = 1.5. Navigator “A” does not have sweeping behaviour (*δ*_2_ is irrelevant in this case). We summarize the key parameters of the navigators in Table 2.

**Table 1.** Model variables and parameters. Let us first note the different notations and their meanings: *A* - constant average wind speed, *ω* - angular rate of change of the wind vector direction, *R* - transverse diffusion of puffs, *f*_*r*_ - puffs release rate, a number of puffs released by the source per second, *r*_0_ - initial radius of a puff at the source, *σ*-rate of growth of a puff, *C*_0_ - odor detection threshold of a navigator,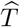 - time it would take a navigator to reach the source along the straight path from the initial position.

| Parameter | Values | Units |
| --- | --- | --- |
| $A$ | 1 | m/sec |
| $B$ | 0 - 0.15 | m/sec |
| $\omega$ | 0.1 | rad/sec |
| $\beta$ | 0 | m/sec |
| $R$ | 0.5 | m<sup>2</sup>/s |
| $f_r$ | 50 – 200 | 1/s |
| $r_0$ | 0.001 | m |
| $\sigma$ | 0.002 | m<sup>2</sup>/s |
| $C_0$ | 500 – 2400 | a.u. |
| $S$ | 200 | pix/s |
| $\hat{T}$ | 1.8-2.4 | sec |

**Table 2.** Key parameters of the navigators simulated in this study. For navigators “C/D”, λ, *δ*_1_, are predetermined constants.

| Navigator | surging | casting | parameter | sweeping |
| --- | --- | --- | --- | --- |
| | $\lambda$ | $\delta_1$ | $\alpha$ | $\delta_2$ |
| A | $t_c$ | $\alpha t_c$ | 1.5 | none |
| B | $t_c$ | $\alpha t_c$ | $\alpha(t)$ | none |
| C | $\lambda$ | $\delta_1$ | 1 | $7\delta_1$ |
| D | $\lambda$ | $\delta_1$ | 1 | $3\delta_1$ |

**Fig 2.**
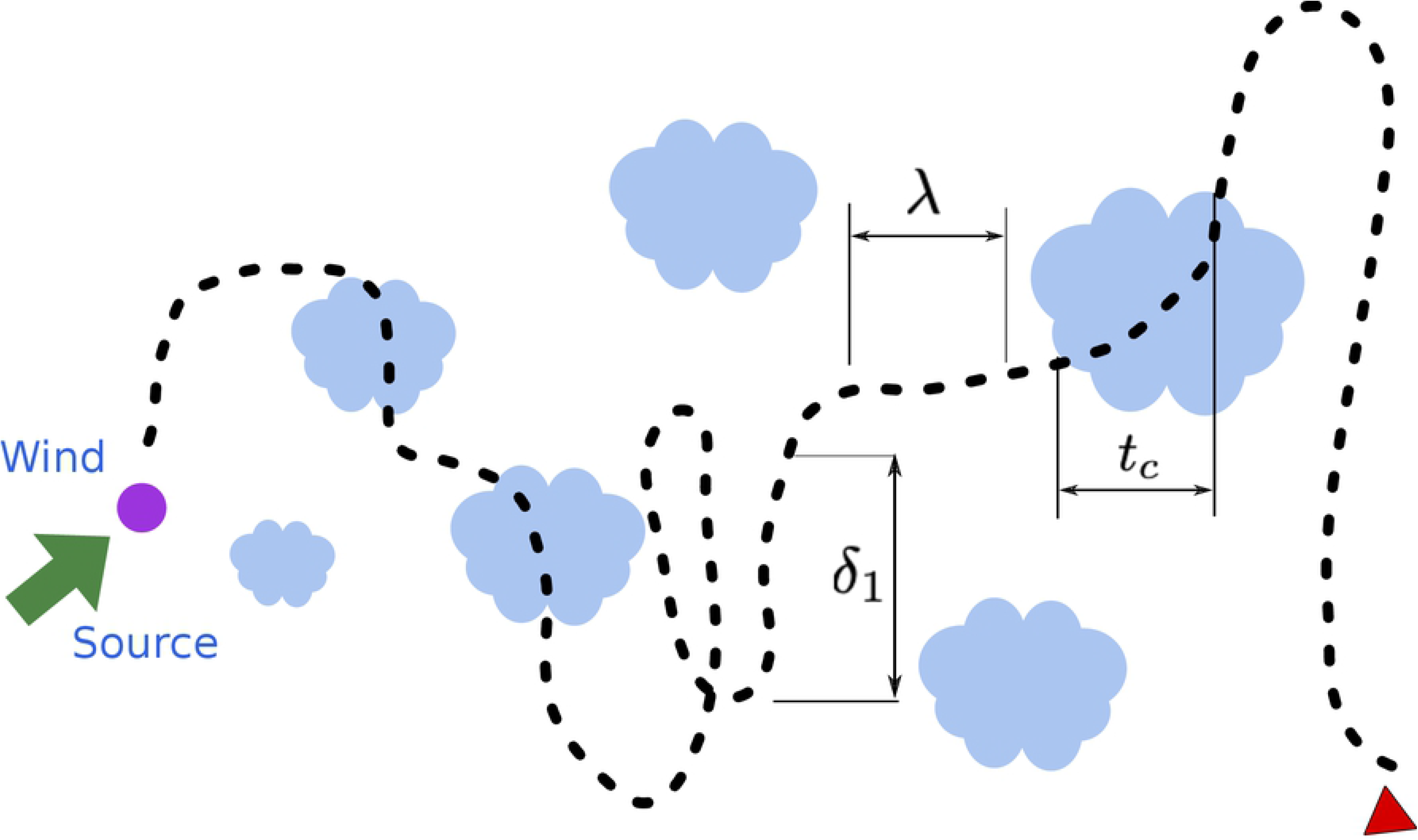
Schematic description of a typical simulation run with the “mean wind” direction from left to right and a typical flight from right to left, including: the two dimensional wind field with an arrow emphasizing an instantaneous direction of wind, a position of the source, a plume of previously released pheromone puffs, and a prototype of a navigation path (type A), following [3]. At the time of detection of the odor, the time of crossing is defined as *t*_*c*_ and used for the following surging (upwind flight for time denoted as λ) and casting (crosswind flight, a typical time denoted as *delta*_1_). Due to the turbulent nature of the odor spreading and diffusion, pheromone puffs are increasing in size with the distance from the source.

#### Strategy “B”

Another navigation strategy which uses the detection/crossing time interval *t*_*c*_ as a basic parameter is called “B”. Effectively it is a small modification of the strategy “A”. In strategy “B”, casting time increases with every other turn, slowly growing and covering a larger cross-wind width. Mathematically we define it as *δ*_1_ = *α*(*t*)*t*_*c*_, where *α*(*t*) marks a continuously growing function with a predefined coefficient. A typical flight path of the navigator type “B” would appear as shown schematically in Fig. 3.

**Fig 3.**
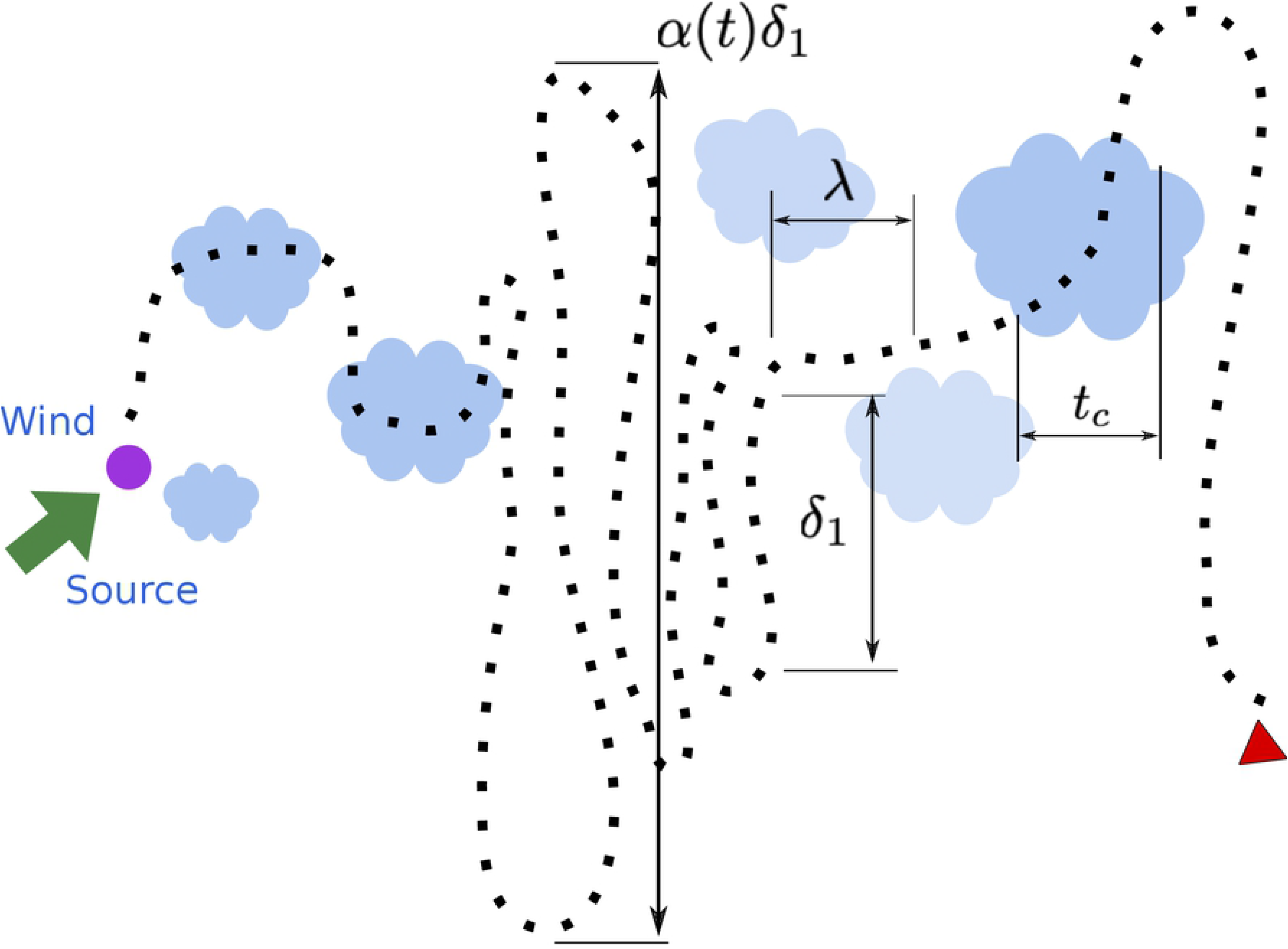
Schematic description of navigator “B” path - in addition to the cross-wind casting option, this navigator type increases its following casting time (or width of search) at some constant rate, i.e. *δ*_1_ = *α*(*t*)*t*_*c*_, where *α*(*t*) is a predefined, non-adaptive, rate.

#### Strategies “C” and “D”

Navigation strategies “C” and “D” are based on the ideas in Ref. [2]. The basic move is that anemotaxis, i.e. after every detection, the navigator surges upwind for time λ. When odor is lost, i.e. there are no new detections after a certain predefined time interval λ, the navigator changes to the casting mode with another predefined time interval *δ*_1_. After several turns (an arbitrary suggested number of turns that can be defined following the literature, here it is set to 7), the navigator will perform a large sweep, as shown in Fig. 4. The time interval which characterises the sweeps, *δ*_2_, varies between strategies “C” and “D”. For strategy “C” *δ*_2_ = 7*δ*_1_, while for strategy “D” *δ*_2_ = 3*δ*_1_. This parameter can basically differentiate the success rate of flyers with large sweeps versus small sweeps. Note that there is a hidden parameter in the sweeping behaviour and it is a randomly chosen angle with respect to the wind. Straightforwardly speaking it can be a “good choice” and the sweep will cross the centerline of the simulation field and increase the probability to detect another puff, or a “bad choice” (with 50/50 probability according to the random decision of the angle) that will distant a navigator from the plume region, such that only another randomly chosen large sweep in the opposite direction can bring the moth back closer to the plume. In some cases, sweeps will move the navigator outside of a simulation boundaries, effectively stopping the simulation for the given navigator.

**Fig 4.**
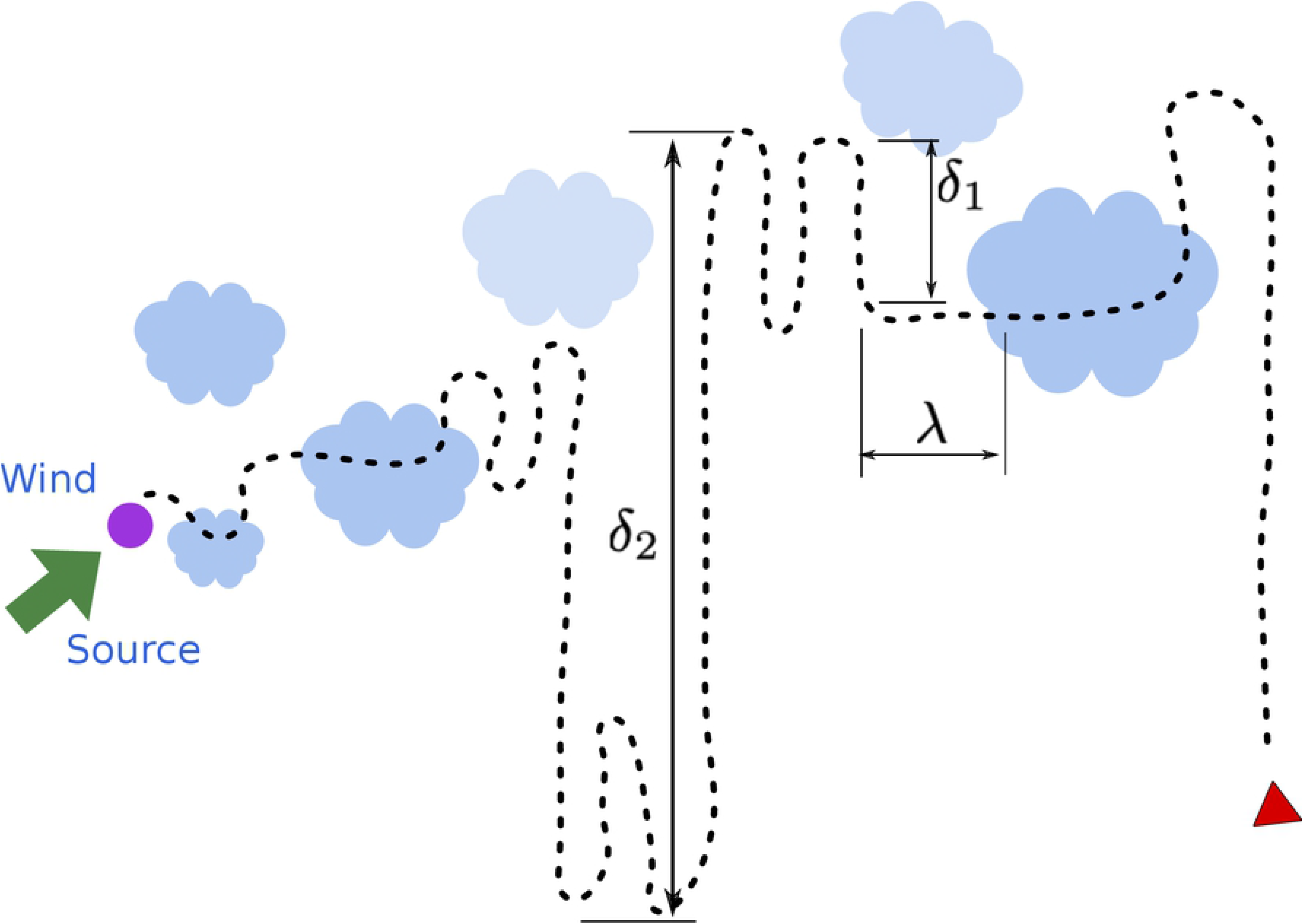
Schematic description of navigators “C” and “D”, both have an additional navigational behaviour of a “sweep” at a different appearance rate and at different amplitude - large motion in a cross-wind direction at time scale *δ*_2_, alternating with the casting behaviour (small zigzags) at the time scale *δ*_1_.

The strategies were designed for the sake of comparison and only inspired by the work of Ref. [2] and the references therein. This study is not focused on selecting a better or optimized navigation strategy or suggests the best solution. We compare the navigators as a case study to demonstrate the strength of the simulation platform and its robustness to model various strategies. To summarize, we compare two navigation strategies with the adaptive parameters, depending on the last puff detection and the pre-determined strategies that are based on a constant set of parameters. Furthermore, we test for any effect due to increasing casting width with the time of search and test whether small/large sweeps could be a mechanism that resets the navigator to the centreline and renews a successful anemotaxis approach. The effects of the two additional features (increasing casting width and sweeping) are especially interesting with respect to the meandering of the plume. The modeled plumes can propagate along very curved centrelines during the moth flights, hindering the anemotaxis concept.

## Results

Using the aforementioned methods and software, we created a simulator that can mimic different plumes: laminar or turbulent, continuous or sparse, patchy plumes, and strong winds that can have strong gusts or meandering. Comparing different strategies and characterizing their sensitivity to the plume, wind field and navigator parameters, can assist in addressing the key ingredients of an olfactory-like navigation. In the following, we present results of the simulator in which we compare the strategies’ sensitivity to two parameters: meandering and a puff spread rate.

We present randomly chosen “flights” or navigation paths by navigators “A-D” in Fig. 5. The flights are drawn as continuous curves, although we simulated them as discrete points at fixed time intervals. We plot the results according to the coordinate system of the simulation: the wind is moving from left to right (increasing *x*) and therefore moths move from right to left (decreasing *x* towards *x* = 0). A successful navigation trajectory ends in the proximity of the origin *x* = 0, *y* = 0. The randomly chosen flights in Fig. 5 do not imply that some are more or less successful, they merely demonstrate a shape of a single navigator from the large population released during a simulation run. We compare statistics of all the released navigators moving in the same simulation field. Then the simulation field is updated with the new parameters (e.g. increased meandering) and multiple navigators are released again. To equalize the conditions, each group of navigators is placed at the same starting points on a rectangular grid of 200 columns and 60 rows, totaling in 12,000 navigators for each strategy. The navigators are independent and cannot interact with each other.

**Fig 5.**
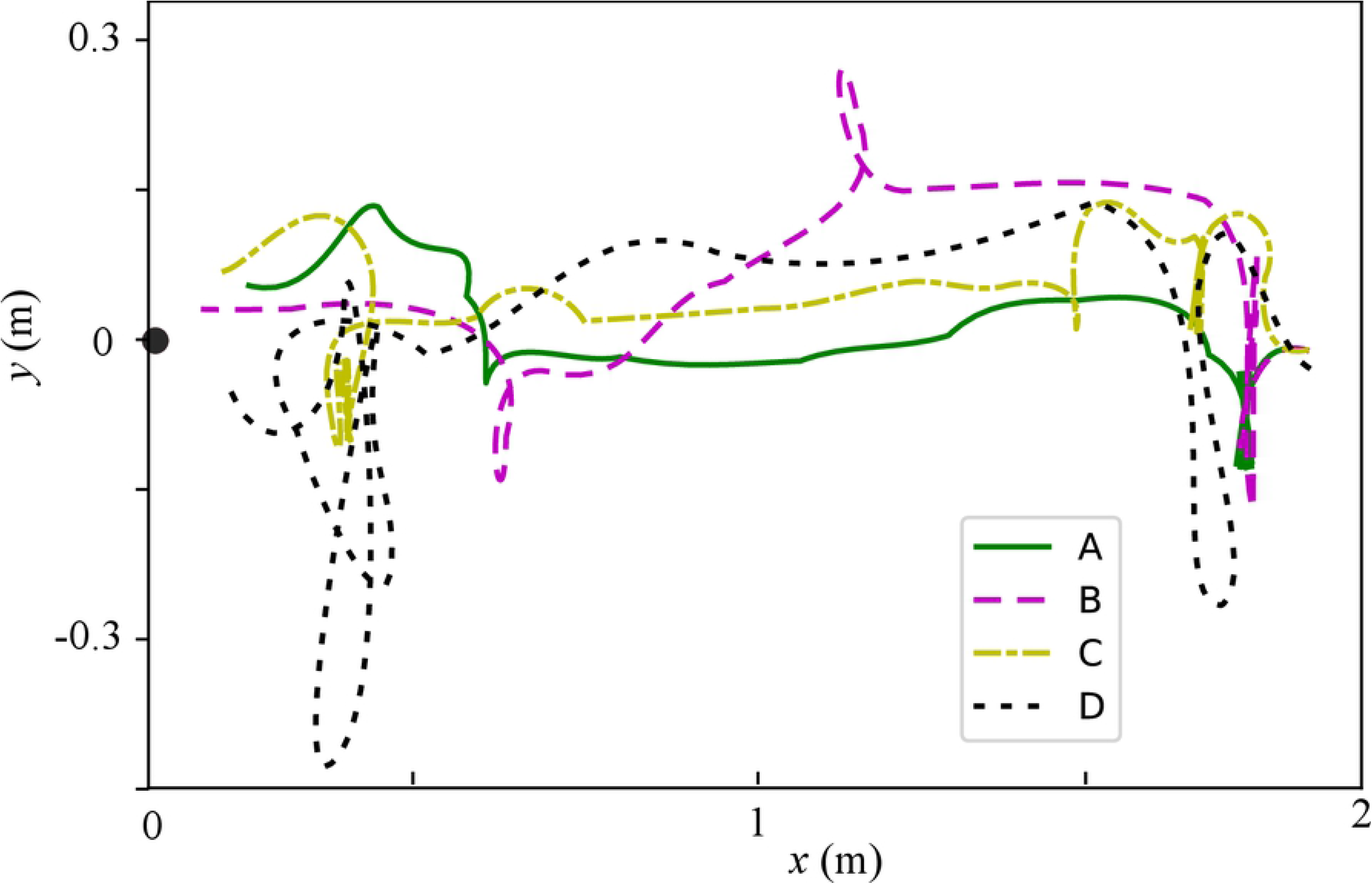
Typical flight paths of the four navigators strategies “A-D”. The paths started at large *x*_0_ (on the right side of the simulation field) and navigated towards the origin *x* = 0. The units are meters. All the navigators have the same ground speed.

### Sensitivity of strategies to wind and plume parameters

All the four strategies “A-D” are capable of successful flights in present simulations. Our focus is not on the extreme cases, where only a specific navigation strategy may have an advantage, but rather on a statistically significant set of flights that will help us find the sensitivity of the navigation strategies to some key parameters. The first key parameter is the meandering of the plume. As reported in the previous studies, moving upwind (anemotaxis) is a well thought strategy, but theoretically cannot work in a strongly meandering wind. For instance, if the navigator is using local wind direction at the time it encounters the first puff, during this period of time of the puff propagation, the wind completely changes its direction - then the navigator will receive an unreliable signal on the source location.

The ability of the navigators to find the plume source is classified using the two metrics:

- Success rate - is the percentage of navigators that reached the origin (within a short distance of 0.15 m) out of all moving navigators (navigators which did not encounter any odor and consequently did not begin their search were omitted from this calculation). A higher percentage for a specific navigation strategy, as compared to other strategies (for the same wind and plume parameters) could imply a more successful navigation strategy. Of course the time of flight, the number of search maneuvers and other parameters should be taken into account for more meaningful comparison.
- Average navigation time, *τ* (ratio) - the average ratio between the time of navigation and the minimal theoretical navigation time, as expressed in the following equation, 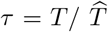. Only successful navigation paths are taken into account in this calculation, it is a measure of the navigation efficiency: smaller ratio means a more efficient navigation algorithm.

### Sensitivity to the meandering amplitude

In this subsection we present the ability of different navigators to reach the odor source in different meandering conditions. Meandering determines the extent in which the wind changes direction between the puff release, first puff arrival to the flyer’s initial position and the navigation process. Meandering is characterized here by two parameters: amplitude and time period, as defined in Eq. (1). A stronger amplitude and shorter period will also affect the separation of puffs within the plume, in addition to creating a“curved” or “wavy” plume shape. We keep the time period constant and vary only the meandering amplitude, the parameter shown on the horizontal axis in Fig. 6.

**Fig 6.**
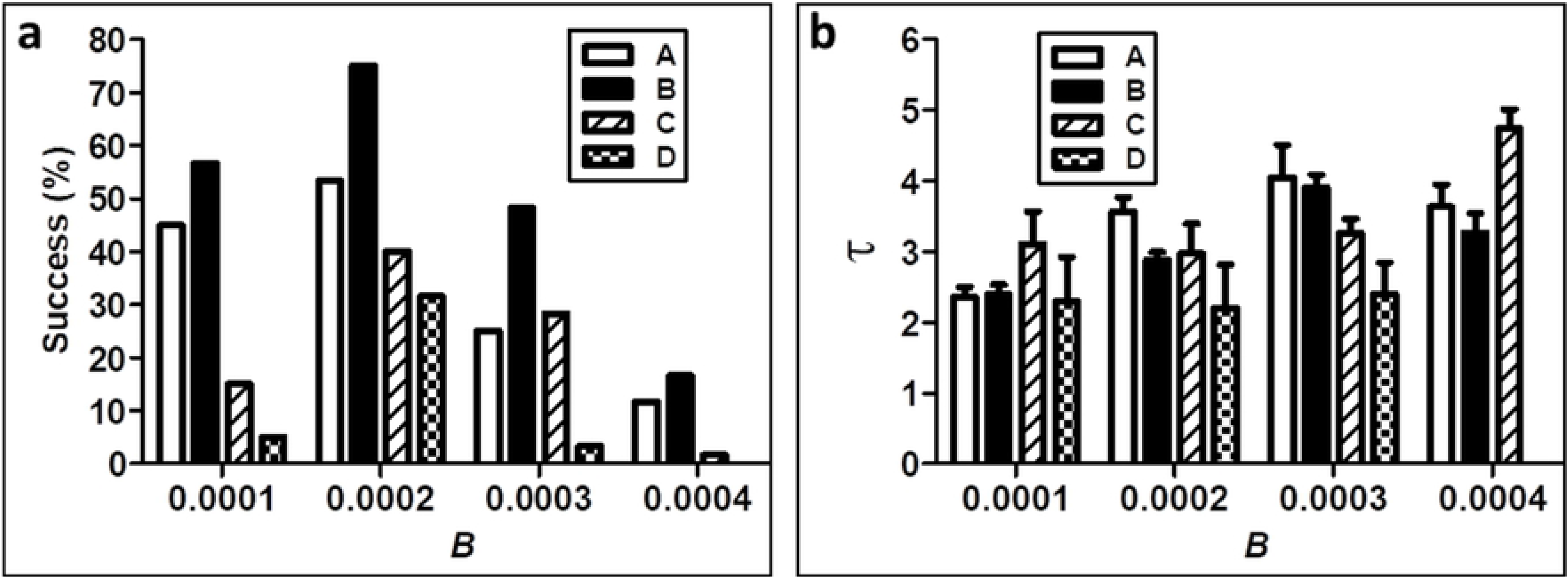
Comparison of different navigation strategies in increasing wind meandering amplitudes in terms of: a) the success rate, and b) flight time. *τ* is the relation between the mean flight time of a single navigator group and the theoretically best possible flight time (a straight line from the starting to finishing point). The lower the value of that relation, the faster (“better”) the navigation. Each column presents the mean result for a group of navigators of the same type.

Fig. 6 presents the statistics of navigators of types “A-D”, flying in the same domain in which we vary only the meandering amplitude between simulation runs. For all navigators, the success rate decreases with increasing amplitude of meandering (beyond the first jump from 0 to 0.05). The average time does not change significantly with the increasing meandering amplitude, however, this result should be seen with respect to the much lower number of successful flights. Note that this statistic is only calculated for successful flights. Some columns disappeared because of the navigators that do not complete their navigation successfully for large meandering amplitudes. Apparently, the adaptive navigation strategies have some advantage for the stronger meandering plumes. That is plausible due to their ability to change the search depending on the previous detection time.

### Sensitivity to the spread rate

The rate of puff spread in real flows is a complex function of the turbulent flow field that can be formulated as shown in Eq. (3). In order to mimic the increase in turbulence intensity (level) during the simulations in a simplified manner, we increase the radii of puffs. For a navigator, increasing turbulence intensity could infer, on one hand, a higher probability of finding a puff, but on the other hand, the entire puff may be below the detection rate, due to a high level of mixing at shorter distances from the source. In Fig 7 we visualize a possible apparent effect of turbulent intensity on the simulation fields of the same plume parameters with different puff spread rates.

**Fig 7.**
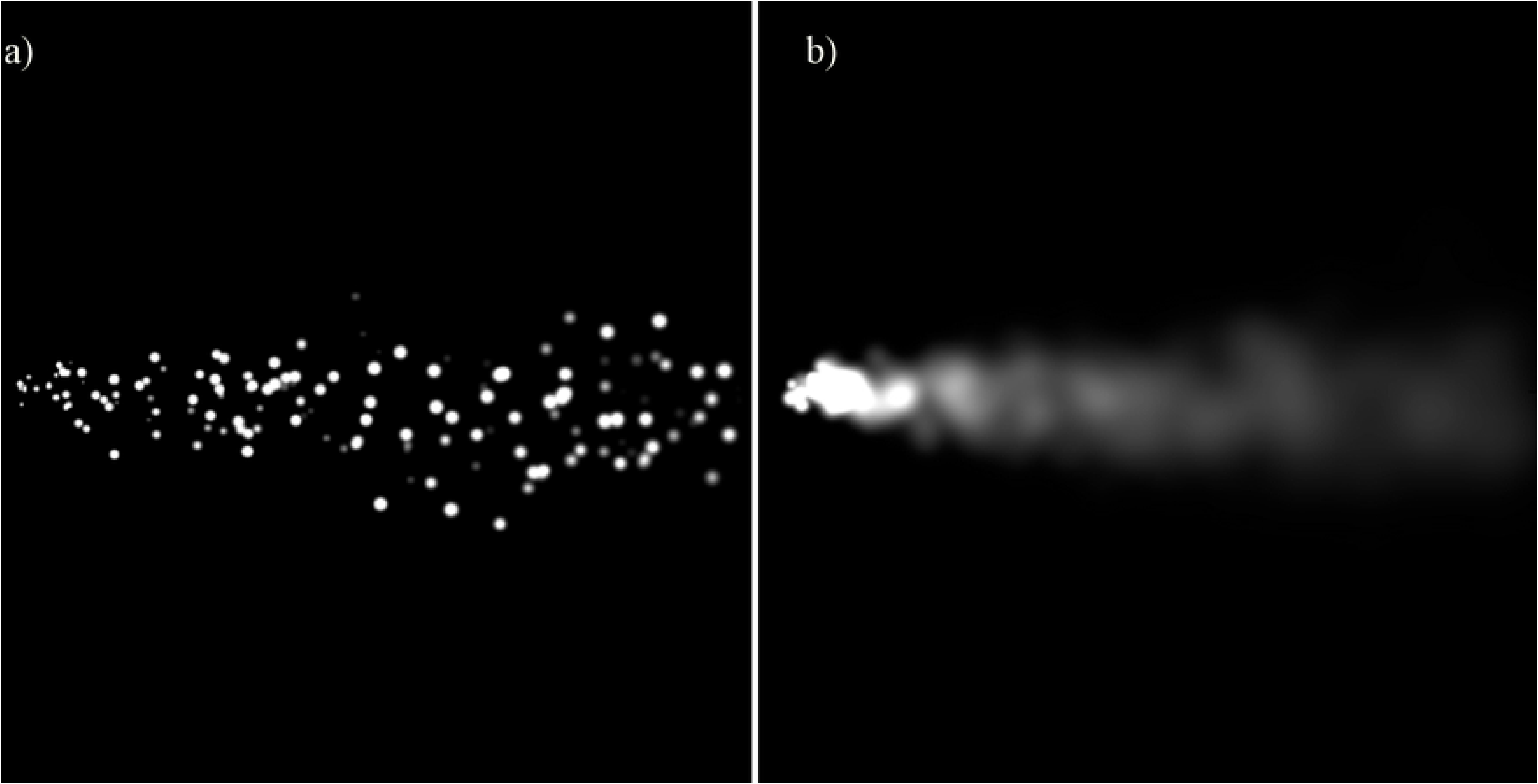
The turbulent plume as it appears to the navigator for the different levels of the spread rate, see Table 2. An increase in puff spread rate effectively simulates an increase in turbulent intensity. a) puff spread rate is 0.0005 m^2^/s, creating a stream of distinct, separate puffs; and b) puff spread rate is 0.001 m^2^/s, leading to overlap between different puffs that results in a continuous plume.

In Fig 8 we compare the different navigation strategies for increasing turbulent intensity. Note that the relation between average navigation time as well as success percentage are not monotonic. For a given set of conditions, it appears that there is an “optimal” turbulence intensity level, and the rest of the values change for increasing/decreasing values for different reasons. In this particular case, there is a positive effect of increasing search interval *δ*_1_ is evident in results of navigator “B” in terms of the success rate and time, as compared to type “A”.

**Fig 8.**
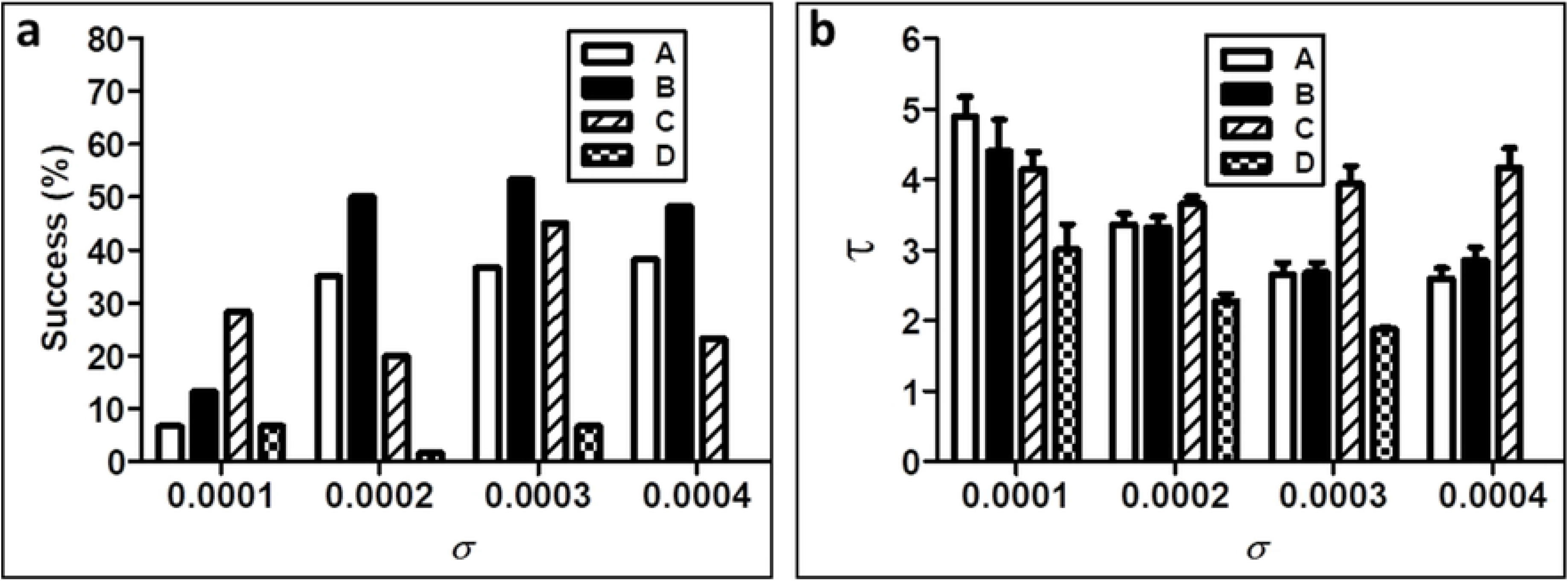
A comparison of different navigation strategies in turbulent conditions, as determined by the puff spread rate.

One plausible explanation is the strict dependence of type “A” on the last measurement of the crossing time which becomes a less reliable signal for high spread rates and especially at small distances from the source. Navigator types “C” and “D” seem to respond negatively to the increasing spread rate, though similarly navigator “C” (Large *δ*2) outperforms navigator “D” (small *δ*2).

## Summary and discussion

The main goal of this work is to create a computational framework for comparison of various navigation and decision making strategies. It is not necessarily limited to flying objects, with small modifications it could also fit to simulate walking or swimming agents. It is designed to allow comparison of strategies also for diffusive spread of scalars, with zero wind speed, as well as problems with multiple sources.

In this case study we implemented four navigation strategies designed to track a chemical plume trail, created by a pulsating source in a turbulent flow. The navigators are self-propelled flyers with a single sensor that provides a binary odor detection (above/below a threshold), and a timer, but without memory of previous states. Elaborating on an existing computational simulation of turbulent flow [33], we propose a system to simulate and evaluate the performance of such navigation algorithms. We present a case study for 12,000 navigators of four different types and multiple parameter changes of wind and plume parameters. Each navigator can only access the data about the pixel on which it is positioned - the local wind velocity (magnitude and direction), and local odor concentration below/above its sensor threshold. Comparison of different strategies is performed statistically initiating a large set of randomly varying parameters within a given set of constraints and conditions. For each instance, we locate a group of navigators and collect statistics of this population in a sort of Monte Carlo simulation. The important constraint is to use the same predefined simulated wind and plume fields, as well as the same starting positions for a group.

In this paper we focused on strategies using fixed parameters [2] (“C/D”), with the strategies that use locally available information about the puffs, as suggested in [3] (“A”), and augmented with the extended casting search type “B”. For the sake of comparison, a large set of runs was performed and the results are compared in terms of efficiency parameters: shorter/longer average navigation time, and a success rate (the number of successful searches out of all moving navigators). We provide the comparison in terms of histograms and compare our navigators for sensitivity to various wind or plume characteristics. It is not the purpose of this work to claim that one of the types has a better or worse performance, as this statement would be valid only in a context of the simulated parameters. We observe that some navigation strategies could imply an adaption to strongly meandering winds, which could potentially be an important insight comparing flights of insects from different regions, or habitats.

The future research that will be based on the developed framework could assist development of better navigating strategies for autonomous drones or underwater vehicles, e.g. [31] or analysing sophisticated navigation strategies such as infotaxis [28], [27].

The presented simulation framework is not developed with the real physical models in hand. For instance, the puff spreading rate which is proportional to the turbulent intensity in atmospheric boundary layers is taken here as a constant, despite the fact that it is a very complex function of a varying turbulence level, a proximity to the trees and canopies, or strongly depending on the atmospheric boundary layer stability [37]. In the future, it would be possible to extend this study with more realistic fluid dynamics and scalar dispersion models. It would be also valuable to extend this work to a realistic 3D simulation field that could take into account also height varying parameters and a possible effect on foraging and mating insects.

## Acknowledgements

This study is supported by the U.S.-Israel Binational Science Foundation (BSF) under grant 2013399 and the Ministry of Agriculture under grant 13-28-0003.

